# The AmiC/NlpD pathway dominates peptidoglycan breakdown in *Neisseria meningitidis* and affects cell separation, NOD1 agonist production, and infection

**DOI:** 10.1101/2021.09.02.458811

**Authors:** Jia Mun Chan, Kathleen T. Hackett, Katelynn L. Woodhams, Ryan E. Schaub, Joseph P. Dillard

## Abstract

The human-restricted pathogen *Neisseria meningitidis*, which is best known for causing invasive meningococcal disease, has a nonpathogenic lifestyle as an asymptomatic colonizer of the human naso- and oropharyngeal space. *N. meningitidis* releases small peptidoglycan (PG) fragments during growth. It was demonstrated previously that *N. meningitidis* releases low levels of tripeptide PG monomer, which is an inflammatory molecule recognized by the human intracellular innate immune receptor NOD1. In this present study, we demonstrated that *N. meningitidis* released more PG-derived peptides compared to PG monomers. Using a reporter cell line overexpressing human NOD1, we showed that *N. meningitidis* activates NOD1 using PG-derived peptides. Generation of such peptides required the presence of the periplasmic *N-*acetylmuramyl-L-alanine amidase AmiC, and the outer membrane lipoprotein, NlpD. AmiC and NlpD were found to function in cell separation, and mutation of either *amiC* or *nlpD* resulted in large clumps of unseparated *N. meningitidis* cells instead of the characteristic diplococci. Using stochastic optical reconstruction microscopy, we demonstrated that FLAG epitope-tagged NlpD localized to the septum, while similarly-tagged AmiC was found at the septum in some diplococci but distributed around the cell in most cases. In a human whole blood infection assay, an *nlpD* mutant was severely attenuated and showed particular sensitivity to complement. Thus, in *N. meningitidis* the cell separation proteins AmiC and NlpD are necessary for NOD1 stimulation and for survival during infection of human blood.

## INTRODUCTION

*Neisseria meningitidis*, also known as the meningococcus, is an obligate colonizer of the human nasopharyngeal space. Meningococci occasionally disseminate to cause invasive disease such as meningitis and septicemia. Invasive disease has a fatality rate around 11% even with early treatment, and 11-19% of survivors develop lifelong sequelae [1-3]. *N. meningitidis* colonizes up to 25% of the population at any one time [3]. The carriage rate of *N. meningitidis* can be as high as 70% in areas of high density such as military barracks, college dormitories, and during the Hajj pilgrimage [3, 4]; prolonged exposure to carriers increases the risk of invasive meningococcal disease for non-carriers [4]. *N. meningitidis* encodes multiple virulence factors that allow it to invade the bloodstream and cross the blood-brain barrier, cause a large inflammatory response, and evade clearance by the immune system [5, 6]. Much of the tissue damage sustained during meningococcal disease is a result of a strong inflammatory immune response [3, 6, 7]. One class of immunostimulatory molecules made and released by *N. meningitidis* is diaminopimelic acid (DAP)-type peptidoglycan fragments [8].

Peptidoglycan (PG) is a structural macromolecule that confers bacterial cell shape and protects the cell against turgor pressure. It is made of repeating subunits of *N-*acetylglucosamine (GlcNAc) and *N-*acetylmuramic acid (MurNAc), with peptide chains attached to the *N-*acetylmuramic acid moiety [9]. The third amino acid on this peptide chain is DAP in most Gram-negative bacteria and Gram-positive bacilli, and lysine in most other Gram-positive bacteria [10]. Breakdown of existing PG strands and incorporation of newly synthesized PG strands allow for cell enlargement and cell separation [11], and this process generates small PG fragments as the bacteria grow [12, 13]. Most Gram-negative bacteria release very small amounts of PG fragments; instead, PG fragments are typically contained in the periplasmic space and efficiently taken back into the cytoplasm by a PG fragment permease in order to be reused in cellular processes [13, 14]. A limited group of Gram-negative bacteria, including but not limited to *N. meningitidis*, the closely related pathogen *N. gonorrhoeae*, the etiological agent of whooping cough *Bordetella pertussis*, and the squid symbiont *Vibrio fischeri* release cytotoxic PG monomers (GlcNAc-anhydroMurNAc-peptide) with no overt effects on bacterial growth [8, 12, 15, 16]. PG monomers released from *N. gonorrhoeae* cause ciliated cell death in human Fallopian tubes [15, 17]. In addition to releasing PG monomers, *N. meningitidis* also releases PG dimers (two glycosidically- or peptide-linked monomers), and PG-derived sugars such as GlcNAc-anhydroMurNAc disaccharide and anhydroMurNAc [8]. The release of PG-derived peptides by *N. meningitidis* has not been characterized.

Host organisms such as humans have evolved strategies to detect and respond to microbe associated molecular patterns to either prevent or resolve an infection. One such intracellular innate immune receptor is human NOD1 (hNOD1). hNOD1 induces a proinflammatory response upon recognition of PG fragments with a terminal DAP, which includes tripeptide PG monomers (GlcNAc-anhydroMurNAc-L-ala-D-glu-*meso*DAP) and PG derived tripeptide (L-ala-D-glu-*meso*DAP) [18-21]. hNOD1 does not respond well to amidated PG fragments; some species of Gram-positive bacteria that have DAP-type PG, such as *Bacillus subtilis*, amidate the DAP moiety of their PG stems [19, 20, 22]. Thus, hNOD1 predominantly detects the presence of Gram-negative bacteria, which includes *N. meningitidis*. In this study, we characterized PG-derived peptides released by *N. meningitidis* and found that meningococci release immunologically relevant amounts of hNOD1 activating PG-derived peptides.

## RESULTS

### The Ltg and the AmiC pathways of sacculus degradation

Previous studies of PG fragment breakdown and release by *N. meningitidis, N. gonorrhoeae*, and commensal species indicate that PG can be broken down in two different ways (Fig. 1) [15]. In the lytic transglycosylase-dominated pathway, LtgA removes peptide-crosslinked dimers from the sacculus, and an endolytic lytic transglycosylase, likely LtgE, creates glycosidically-linked dimers [23, 24]. After DacB (PBP3) cuts crosslinks between strands, LtgA and LtgD degrade strands to PG monomers [8]. LdcA acts on the tetrapeptide PG monomers to convert them to tripeptide PG monomers, thus making them NOD1 agonists [25, 26]. For the AmiC-dependent pathway, strand separation by DacB is followed by a stripping of the peptides from the strands by AmiC, with NlpD activation [27, 28]. AmiC cannot remove the final peptide [27, 29]. LtgC then degrades the nearly-naked glycan strand, creating free disaccharides and tetrasaccharide-peptide [30, 31]. NagZ breaks down the disaccharides into monosaccharides [32], and NlpC is predicted to break down tetrapeptides [33]. Although both pathways are active in all *Neisseria* species studied, the lytic transglycosylase pathway is slightly favored in *N. gonorrhoeae*. PG monomers are the most abundant PG fragments released by gonococci, but free tri- and tetrapeptides are almost as abundant as PG monomers [27, 34]. In *N. meningitidis*, experiments monitoring ^3^H-glucosamine-labeled PG suggest that the AmiC pathway is favored. Free sugars and tetrasaccharide-peptide make up a larger proportion of PG fragments released by *N. meningitidis* than do PG monomers and dimers [8]. If *N. meningitidis* extensively uses the AmiC pathway, then this species may make significant amounts of NOD1 agonist in the form of free tripeptides.

**Figure 1.**
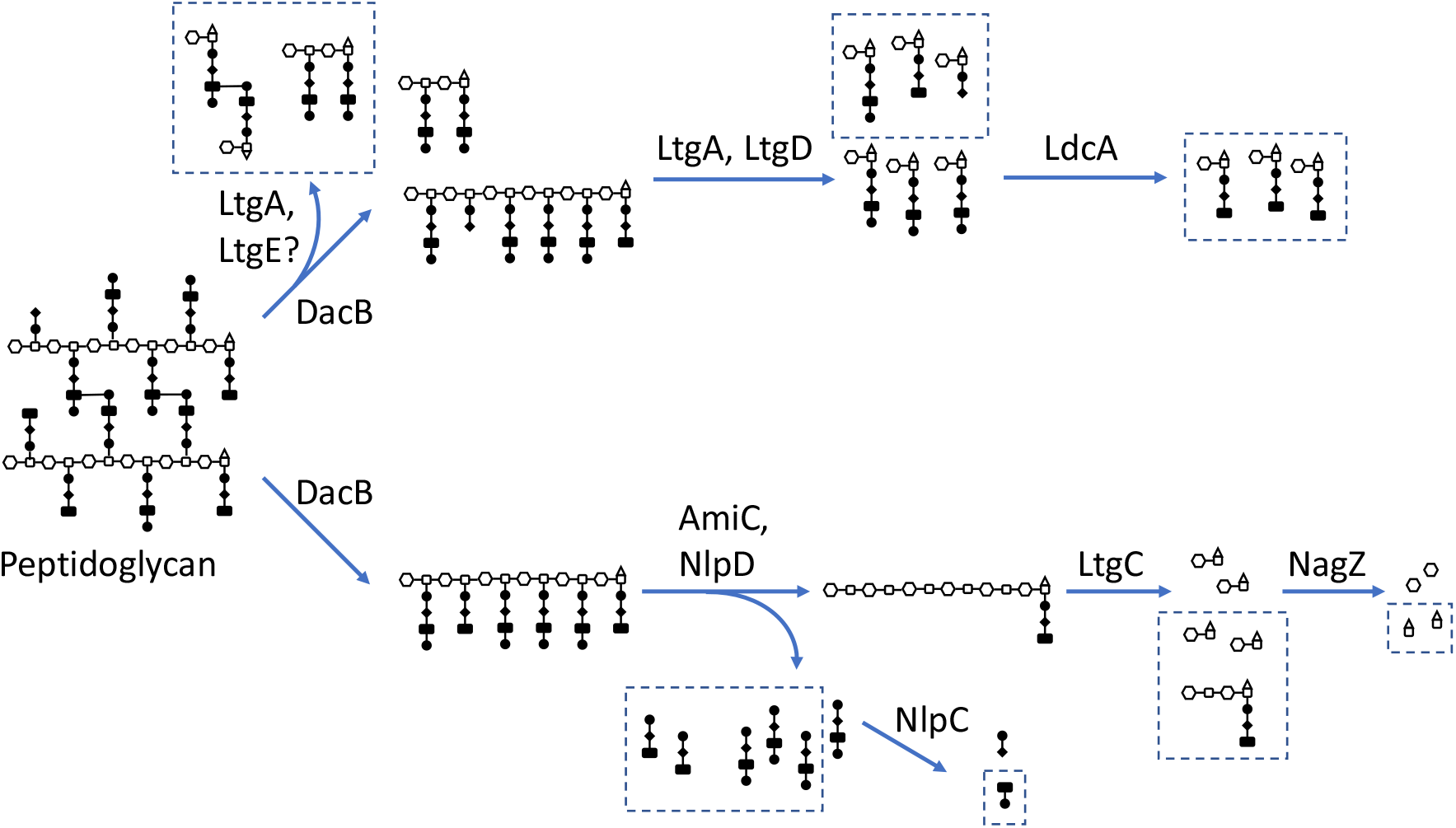
Peptidoglycan degradation pathways in *Neisseria* species. During growth, portions of the sacculus are degraded to allow for cell wall enlargement or remodeling. In the lytic transglycosylase-dominated pathway (top), PG dimers are liberated by LtgA and LtgE, strands are degraded to monomers by LtgA and LtgD, and a majority of PG monomers have their peptides shortened to three amino acids by LdcA. In the amidase-dominated pathway (bottom), peptides are removed from PG strands by AmiC, which is activated by NlpD. Free peptides may be further degraded to dipeptides by one of the three NlpC enzymes. LtgC degrades the denuded strand to free disaccharides and tetrasaccharide-peptide. NagZ, a cytosolic enzyme, cleaves the disaccharides into monosaccharides. Molecules shown with boxes around them are released from the bacteria into the milieu.

We examined PG fragment release in *N. meningitidis* using pulse-chase, metabolic labeling with [2,6-^3^H] DAP. PG fragments released into the medium during growth were analyzed by size-exclusion chromatography (Fig. 2A). Five major peaks were observed, and these contain PG dimers, tetrasaccharide-peptide, PG monomers, free tri- and tetrapeptides, and free dipeptides [8, 34]. The amounts of PG in the free tri- and tetrapeptide and dipeptide fractions were much greater than those in the PG dimer and monomer fractions. These data indicate that *N. meningitidis* extensively degrades its PG to free peptides and sugars prior to release and that it favors the AmiC-dependent pathway for PG breakdown.

**Figure 2.**
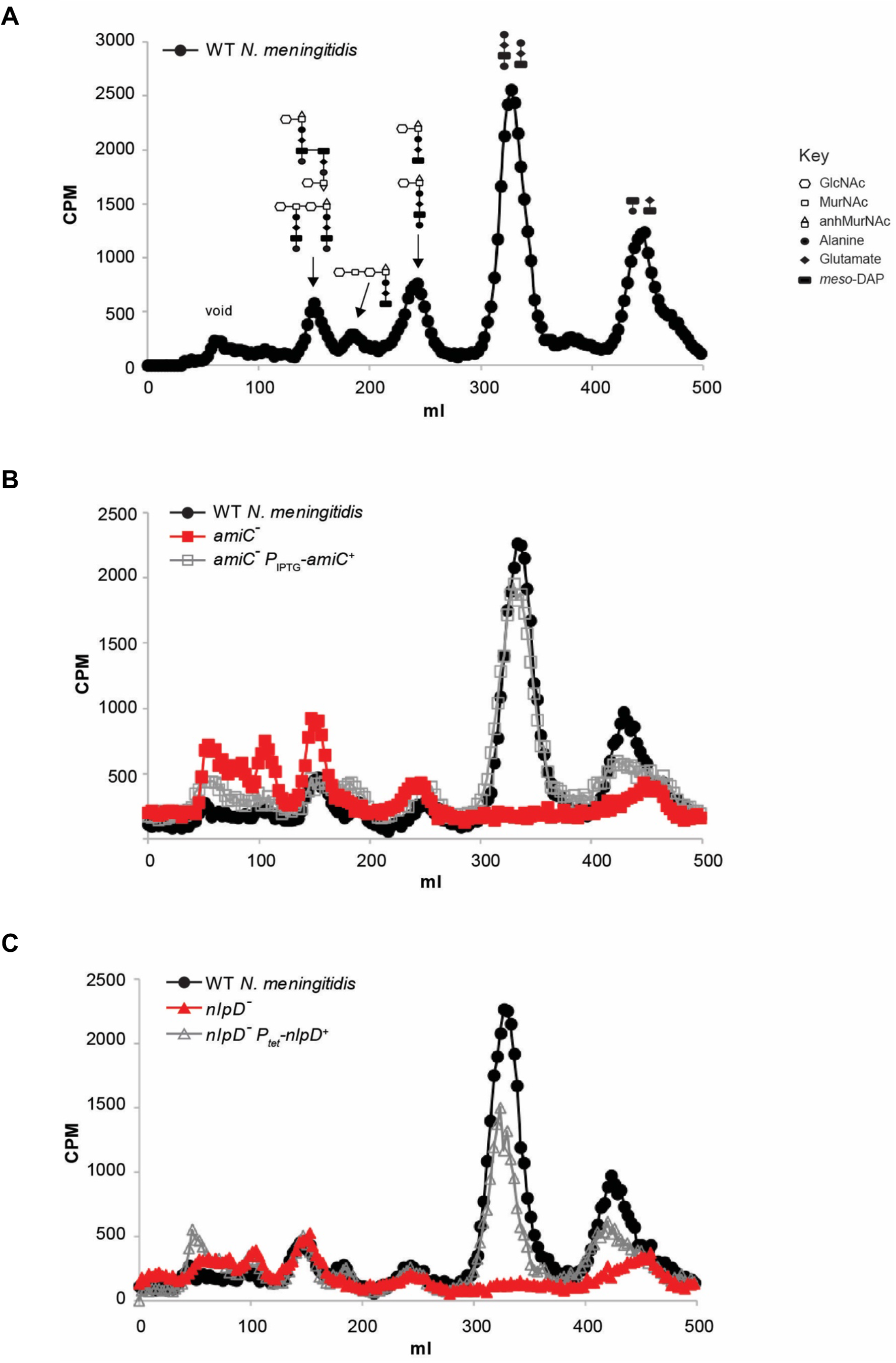
Peptidoglycan fragment release from growing *N. meningitidis*. A) [2,6-^3^H] DAP-labeled PG fragments released by WT *N. meningitidis* (ATCC 13102 *cap*^*-*^) were separated by size-exclusion chromatography, resolving into five distinct peaks in the included volume. Cartoons above each peak show the structures of the PG fragments, representing, in order from the left, dimers, tetrasaccharide-peptide, monomers, free tetra- and tri-peptide, and free dipeptides. Symbols used are based on Jacobs *et al*. (35). The structures of the dipeptides are not yet chemically confirmed. B) PG fragment release from an *amiC* mutant (KL1065) and complement (EC1020). C) PG fragment release from an *nlpD* mutant (KL1072) and complement (EC1026). Mutation of *amiC* (B) or *nlpD* (C) abolished the release of PG-derived peptides. Genetic complementation of the *amiC* (B) and *nlpD* (C) mutations partially restored release of PG-derived peptides. Results depicted are representative of three experiments.

Released tri- and tetrapeptides from PG have been characterized in *N. gonorrhoeae*. Sinha and Rosenthal demonstrated that the peptides were composed of Ala-Glu-DAP and Ala-Glu-DAP-Ala and that they were not crosslinked [34]. Lenz et al. found that an *N. gonorrhoeae* mutant defective for *amiC* did not release free tri- and tetrapeptides [27]. The second PG-derived peptide peak (dipeptides) has been noted previously but not thoroughly investigated, partly because it overlaps somewhat with the peak for free single amino acids not incorporated in the labeling process (Fig 2A), [12, 35]. *Escherichia coli* is known to release the disaccharide DAP-Ala [36]. Therefore, we tested whether metabolic labeling with [^3^H]D-Ala would label the released dipeptide. Incorporation of [^3^H]D-Ala in this peak was similar to incorporation of [^3^H]DAP, indicating that for *N. meningitidis*, as with *E. coli*, the released dipeptide fraction may contain DAP-D-Ala.

### Mutation of *amiC* or *nlpD* abolishes peptide release

*N. meningitidis* encodes one periplasmic *N-*acetylmuramyl-L-alanine amidase, AmiC [37]. Meningococcal AmiC is predicted to be a 417-amino acid zinc-dependent metalloprotease with a Tat signal sequence, an N-terminal AMIN domain that binds PG, and a C-terminal catalytic domain that cleaves peptide stems. AmiC homologues in *E. coli* and *N. gonorrhoeae* function as cell separation amidases [29, 38]. To examine the role of meningococcal AmiC in PG fragment production and release, we generated an in-frame markerless deletion of *amiC*. Mutation of *amiC* abolished PG peptide release and resulted in an increase in PG multimer release (Fig. 2B). If AmiC were able to cleave PG monomers, we would expect to see a proportional increase in PG monomer release with the decrease in peptide release by the *amiC* mutant. However, there is no significant difference in PG monomer release between WT and the *amiC* mutant. Complementation of *amiC* from an ectopic locus restored almost WT distributions of released PG fragments. Our results suggest that meningococcal AmiC liberates peptide stems from the sacculi, and it is necessary for the release of both free tri- and tetrapeptides and free dipetide.

NlpD is a putative AmiC activator protein and functions to potentiate the amidase activity of AmiC in *N. gonorrhoeae* and *E. coli* [27, 28]. Meningococcal NlpD is predicted to be a 415-amino acid outer membrane lipoprotein with two LysM PG binding domains and a degenerate LytM/M23 peptidase domain. Like NlpD homologues from *N. gonorrhoeae* and *E. coli*, only 2/4 of the active site residues in the M23 peptidase domain of meningococcal NlpD are conserved [28, 39]. Analyses in *N. gonorrhoeae* and *E. coli* indicate that their NlpD homologues do not have enzymatic function, but instead act as activator proteins for the amidases [27, 28, 39]. Mutation of *nlpD* in *N. meningitidis* abolished free peptide release without affecting release of PG monomers and dimers (Figure 2C). Complementation of *nlpD* at an ectopic site partially rescued peptide release. We conclude that AmiC and NlpD are indispensable for PG-derived peptide release in *N. meningitidis* and that AmiC activity is dependent on activation by NlpD.

### PG-derived peptides released by *N. meningitidis* stimulate NOD1 activation

HEK293 reporter cell lines overexpressing NOD receptors were used to determine if peptide release by meningococci is necessary for induction of a NOD-dependent response. In this assay, NOD activation is measured by determining the amount of NF-κB controlled secreted alkaline phosphatase (SEAP) detected in the medium via a colorimetric assay. HEK293 cells overexpressing human NOD1 were treated with cell-free supernatant from WT, *amiC* mutant, *nlpD* mutant, or induced cultures of the respective *amiC* or *nlpD* complementation strains (Fig. 3A). As described previously and demonstrated here, supernatant from *N. meningitidis* engenders a NOD1-dependent response (Fig. 3, [8]). We found that supernatant from WT *N. meningitidis* induced 3.6x and 7.5x higher levels of human NOD1 activation compared to an *amiC* or an *nlpD* mutant, respectively (Fig. 3A). Supernatant from *amiC* and *nlpD* complementation strains showed a partial phenotypic complementation, with 2x and 3x higher NOD1 responses compared to supernatant from the mutant strains, respectively (p < 0.05) (Fig. 3A).

**Figure 3.**
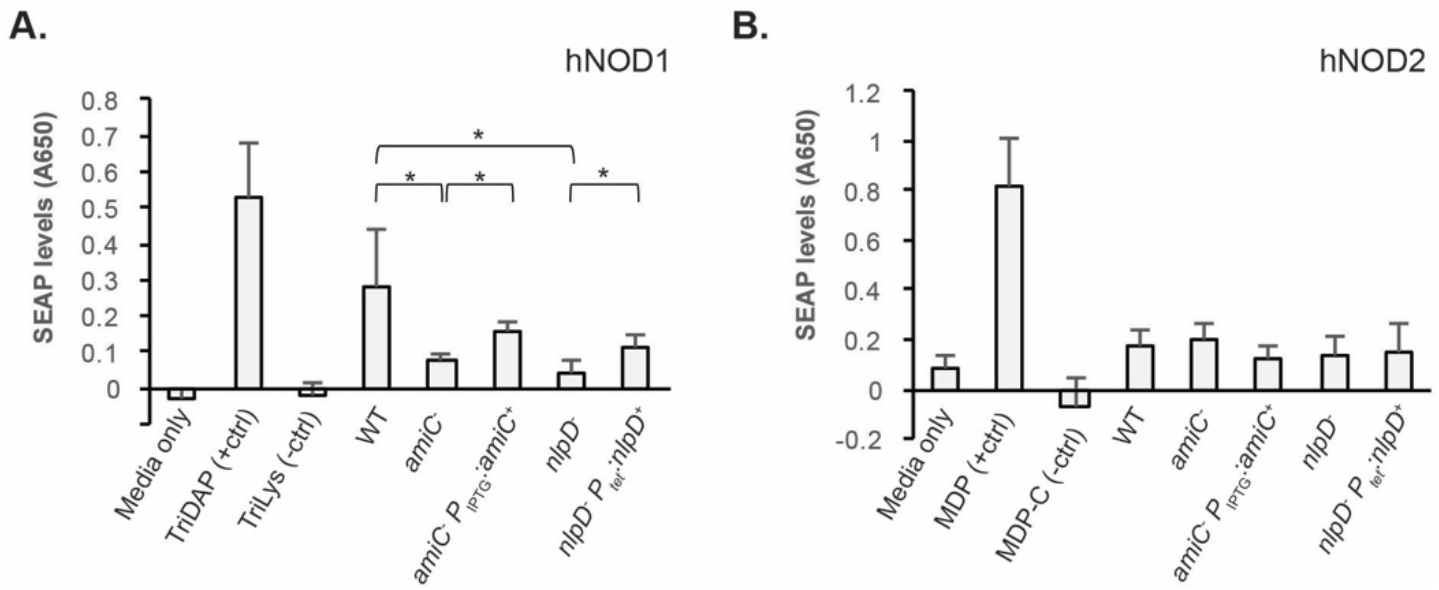
Stimulation of human NOD1 or human NOD2 by conditioned media from *N. meningitidis* mutants that do not release PG-derived peptides. Supernatants from WT (ATCC 13102 *cap*^*-*^), *amiC* mutant (KL1065), *amiC* complementation (EC1020), *nlpD* mutant (KL1072), and *nlpD* complementation (EC1026) strains were harvested, normalized to total cellular protein and used to treat HEK293 reporter cell lines overexpressing hNOD1 (A) or hNOD2 (B), as well as parental HEK293 cell lines. hNOD activation was determined by measuring the amount of secreted alkaline phosphatase (SEAP) in the media. Results are from three independent experiments, with technical triplicates for each biological replicate. Statistical significance was determined by Student’s *t*-test, in which * indicates p<0.05.

We also treated an HEK293 cell line overexpressing human NOD2 in parallel with the same supernatant samples as used for the NOD1 experiments (Fig. 3B). The intracellular innate immune receptor NOD2 recognizes muramyl-dipeptide (MDP) and responds to reducing end PG monomers and thus would not be expected to be differentially affected by amounts of free peptides [40-42]. As predicted, treatment with supernatant from *amiC* or *nlpD* mutants did not alter NOD2 activation levels. Thus, we conclude that PG-derived peptides generated by the action of AmiC and NlpD in *N. meningitidis* induce a NOD1 response, but do not activate NOD2 in a human epithelial cell line.

### AmiC and NlpD are required for cell separation

Periplasmic *N-*acetylmuramyl-L-alanine amidases that act on the sacculi cleave septal PG to allow for proper cell separation [29, 43]. To examine the role of meningococcal AmiC and NlpD in cell separation, we performed thin-section transmission electron microscopy (TEM) to visualize the cell morphology of WT, mutant, and complementation strains. We counted the numbers of cells presenting as monococci, diplococci, tetrads, and 5+ cell clusters in 16-24 fields/strain. Each field used in quantification had an average of 30 cells; an example of a field for each strain used for quantification is shown in Figure 4A. A higher-magnification image demonstrating the size of the multi-cell clumps of the *amiC* and *nlpD* mutants is also included.

**Figure 4.**
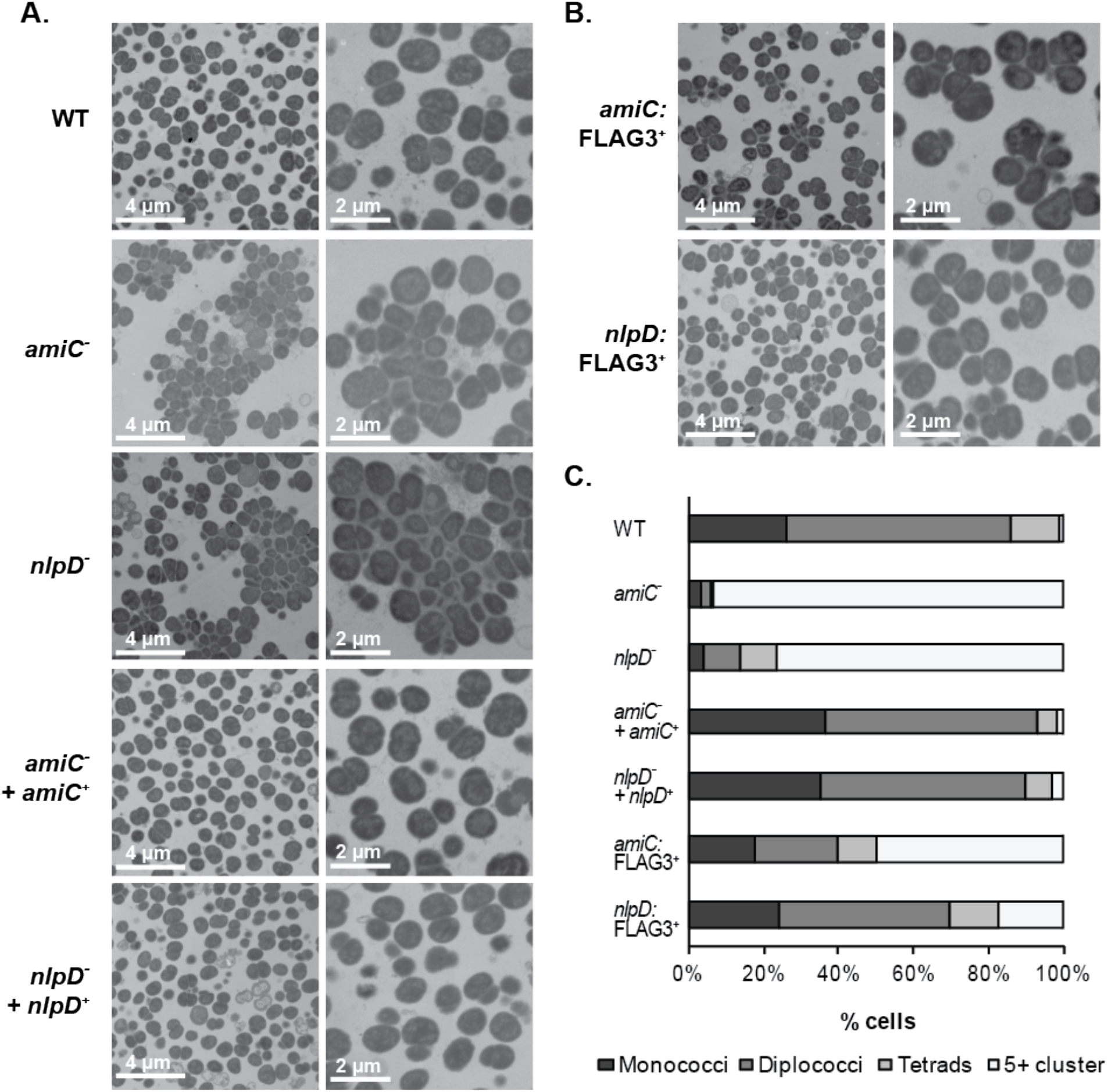
Cell separation defects in *amiC* or *nlpD* mutants. A) WT, *amiC* mutant and complementation strains (KL1065, EC1020), *nlpD* mutant and complementation strains (KL1072, EC1026), as well as B) strains expressing *amiC:FLAG3*^*+*^ (EC1032) and *nlpD:FLAG3*^*+*^ (EC1033) were imaged by thin-section transmission electron microscopy. C) Quantification of the number of cells presenting as monococci, diplococci, tetrads, and clumps of 5+ cells in 16-24 fields of view containing an average of 30 cells each were performed, and presented as percent of total cells.

WT cells predominantly presented as diplococci (60%) and monococci (26%), with few cells found as tetrads (13%) and 5+ cell clusters (1%) (Fig. 4A and 4C). In contrast, both *amiC* and *nlpD* mutants formed large aggregates of 5 cells or more (94% and 76%, respectively). Mutation of *amiC* caused a more severe cell separation defect than did mutation of *nlpD*, as more cells were found in large clusters and fewer cells were found as diplococci or monococci (Fig. 4A and 4C). Only 3% each of *amiC* mutant cells presented as diplococci and monococci, while 10% and 4% of *nlpD* mutants presented as diplococci and monococci, respectively (Fig 4C). Both *amiC* and *nlpD* complementation strains showed WT-like cell morphology, with most cells presenting as diplococci (57% and 55%, respectively) and monococci (37% and 35%), respectively (Fig 4C). We conclude that AmiC and NlpD are required for proper cell separation in *N. meningitidis*.

### C-terminal FLAG3 tagged AmiC displayed both septal and distributed localization while tagged NlpD is septally-localized

We generated C-terminal triple FLAG epitope tagged versions of AmiC and NlpD to visualize localization of these proteins in the cell via stochastic optical reconstruction microscopy (STORM). To determine functionality of these tagged proteins, we performed TEM to look at the cell morphology of the *amiC:FLAG3*^*+*^ and *nlpD:FLAG3*^*+*^ expressing strains (Fig. 4B). The *nlpD:FLAG3*^*+*^ strain predominantly presented as diplococci (46%) and monococci (24%), although it had more 5+ cell clusters compared to WT (17% vs 1%). In contrast, *amiC:FLAG3*^*+*^ cells were more likely to be found in clumps of 5 or more cells (50%) with only 22% and 18% of cells presenting as diplococci and monococci (Fig. 4C). However, the cell clumps in an *amiC:FLAG3*^*+*^ mutant were much smaller and contained fewer cells than the large cell clusters formed by an *amiC* or an *nlpD* mutant, suggesting that AmiC:FLAG3 is partially functional for cell separation.

Localizations of AmiC:FLAG3 and NlpD:FLAG3 in *N. meningitidis* were determined using STORM, and chromosomal DNA was counterstained with DAPI and visualized by epifluorescence microscopy to determine the position of the cells. The presence of a signal band between two adjacent DAPI foci (cells) was interpreted as septal localization. We observed two types of localization patterns for AmiC:FLAG3 (Fig. 5). First, AmiC:FLAG3 localized to the putative septum in a small subset of cells (Fig. 5, left panels). Second, we observed peripheral distribution of AmiC:FLAG3 for most cells in both diplococcal and larger aggregates. NlpD:FLAG3 was clearly localized to the septum in nearly all cells, with a few cells showing some staining that might extend beyond the septum (Fig. 5, right panels). These observations are somewhat similar to the known recruitment of the *E. coli* AmiC and NlpD homologues to the septum during late cell division [38, 44]. Peripheral distribution of AmiC and NlpD homologues in *E. coli* are also seen in non-dividing cells, with a minor enrichment of *E. coli* AmiC at the poles of such cells [38]. Our findings suggest that AmiC:FLAG3 localizes to the septum in dividing meningococcal cells, and may be distributed around the cell in non-dividing cells.

**Figure 5.**
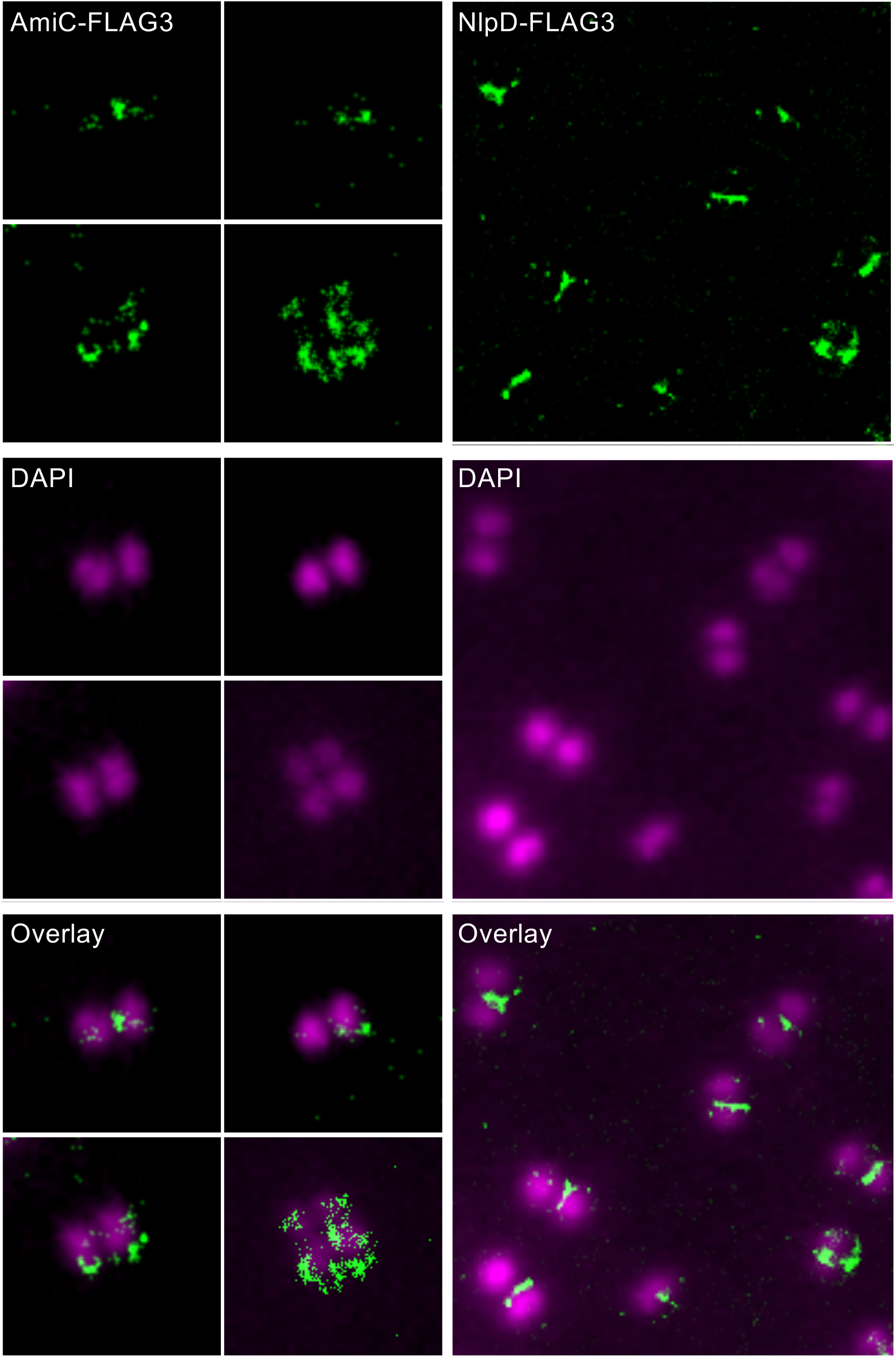
Localization of AmiC:FLAG3 and NlpD:FLAG3 in the meningococcal cell. Localizations of AmiC:FLAG3 or NlpD:FLAG3 were determined using STORM. DNA was counterstained with DAPI. Localization of AmiC:FLAG3 is presented in the left panels, and localization of NlpD:FLAG3 is presented in the right panels. Septal localization of the proteins is seen when they form an approximate line between the two DAPI-stained cells of a diplococcus. As NlpD:FLAG3 exhibited near full functionality, a single field with multiple diplococci was easily found and shown, whereas AmiC:FLAG3 was only partially functional, making it necessary to gather images of diplococci or small clumps from several fields.

### Infection of human whole blood

The severe cell separation defects of the *amiC* and *nlpD* mutants might be expected to affect infection, and differences in the release of PG fragments might affect phagocyte function. However, a significant difficulty in testing the infection ability of such mutants is the inability to accurately quantify the number of viable bacterial cells following infection, since each CFU for the mutants contains multiple bacterial cells stuck together. To circumvent this difficulty, we used the *nlpD* mutant carrying an inducible complement construct. In the absence of the anhydro-tetracycline inducer, the mutant grows as unseparated cell clusters. However, after 4h of growth in medium containing anhydro-tetracycline, a wild-type level of cell separation is achieved (Fig. 6A). Thus we could get a reflection of CFU numbers from infection by growing the bacteria from each timepoint with inducer and then dilution plating the then-separated bacteria.

**Figure 6.**
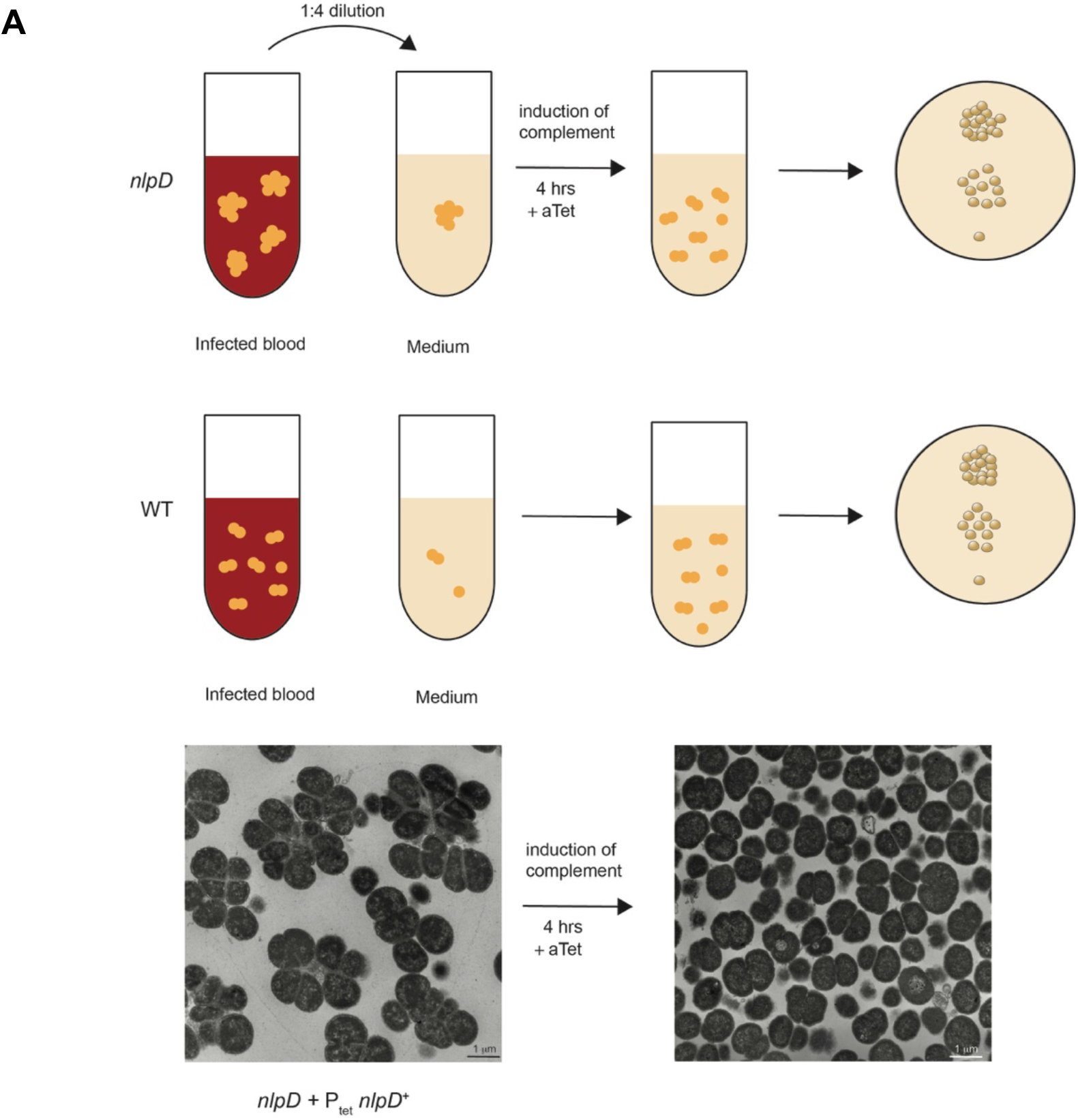
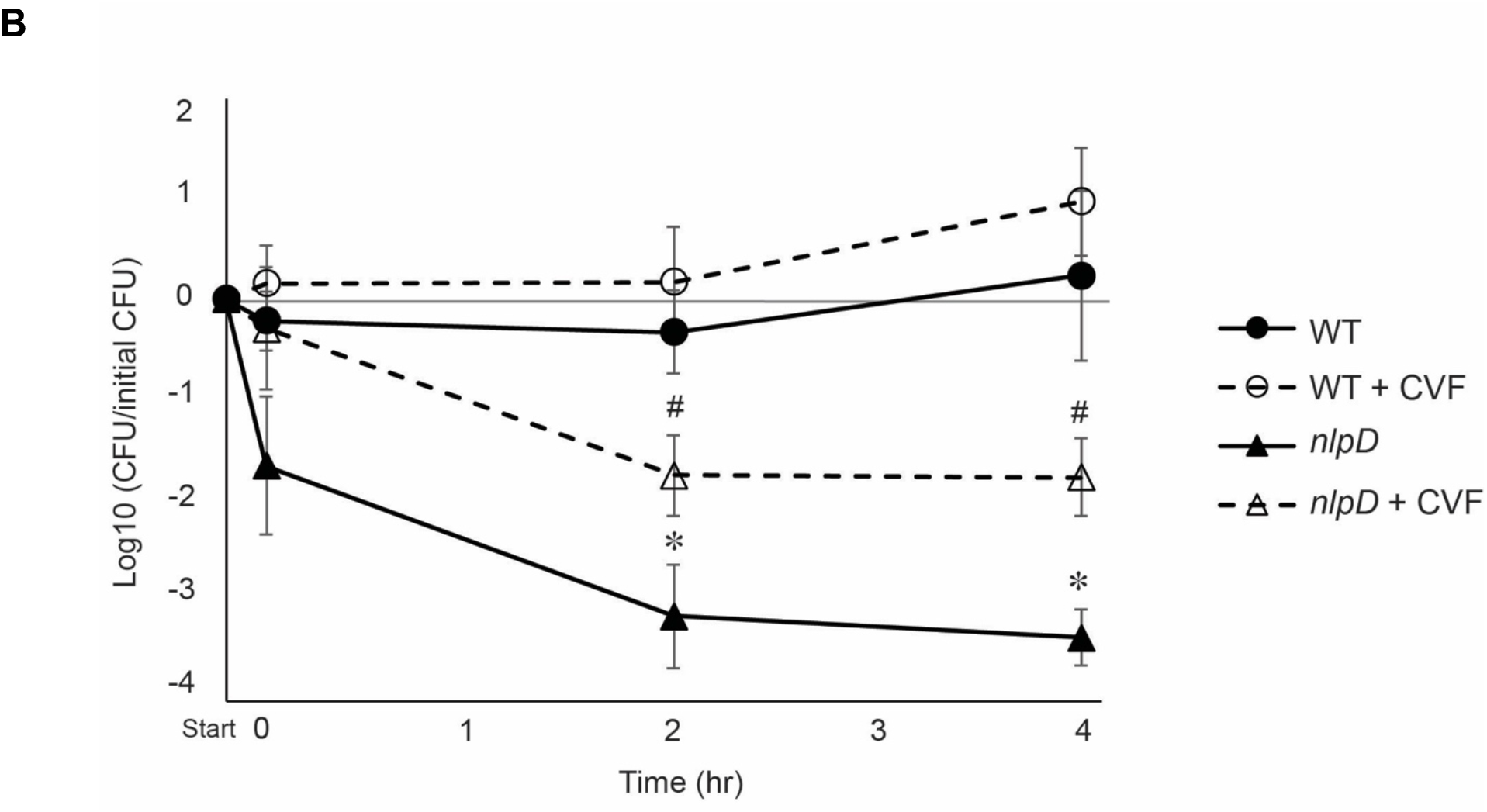
Survival in human whole blood. An *nlpD* mutant with a tightly-regulated complementation construct (EC1026) was used to examine the effect of a cell separation defect on infection. A) Schematic showing the method used for enumerating bacteria following blood infection. Approximately equal numbers of WT and mutant bacteria were used to infect human blood in vitro, though they give different CFU counts due to the *nlpD* mutant not separating. At each time point, the bacteria were diluted into GCBL medium containing anhydrotetracycline to induce *nlpD* expression in the complemented strain, and after 4h of growth the bacteria were diluted and plated to determine CFU numbers. Thin-section transmission electron microscopy shows that EC1026 exhibits the cell separation defect without induction, but 4h in the presence of anhydrotetracycline results in separation to individual diplococci and monococci. B) WT (ATCC13102 *cap*^*-*^) and *nlpD* mutant (EC1026) were inoculated into hirudin-treated human blood. Where indicated, CVF was used to deplete complement in the blood before inoculation. Because of the difficulties inherent in getting equal numbers of separating and not separating bacteria for the initial inoculation, the data were plotted as the ratio of final CFUs to initial CFUs at each time point, on a log scale. Statistical significance was determined by Student’s *t*-test, in which * indicates p<0.05 compared to WT, and # indicates p<0.05 compared to WT + CVF. These data are the geometric mean ± standard error of the mean from five independent experiments.

We compared the ability of the WT strain and the *nlpD* mutant to survive in whole human blood. When we attempted this experiment in an encapsulated strain background, no difference was seen between the WT and mutant. However, in unencapsulated strains, the *nlpD* mutant exhibited a substantial defect. Upon addition of the bacteria to the blood, CFU numbers for the WT strain dropped, reaching ∼50% by 2h and then growing to 170% by 4h (Fig. 6B). By contrast, the *nlpD* mutant dropped 45-fold upon addition to blood and continued to decrease in numbers. By 4h, the *nlpD* mutant had dropped more than 2000-fold. When we used cobra venom factor (CVF) to deplete complement in the blood before bacterial addition, the initial decrease in CFU numbers for the WT strain was eliminated, and the initial decrease for the *nlpD* mutant was reduced to only a two-fold effect. After that point, the WT strain in the CVF-treated blood maintained CFU numbers for the first two hours, then grew to over nine-fold the starting value (Fig. 6B). The *nlpD* mutant in the CVF-treated blood died at the same rate as the *nlpD* mutant did in untreated blood, albeit with CFU values 20-40 fold higher at each time point. Overall, these results demonstrate that the *nlpD* mutant has a defect in survival in human blood and that a large part of that defect is mediated by complement. However, even when complement is depleted, the *nlpD* mutant also shows a survival defect, suggesting increased killing by other soluble factors or phagocytes.

## DISCUSSION

These studies demonstrated that the major PG fragments released by *N. meningitidis* are PG-derived peptides, with the most abundant species being the free tripeptide and tetrapeptide. The tri- and tetrapeptides are also released in large amounts by the related species *N. gonorrhoeae*, though *N. meningitidis* releases much greater amounts of the peptides than *N. gonorrhoeae*. The peptides are generated by the action of the periplasmic *N-*acetylmuramyl-L-alanine amidase AmiC, which is activated by the outer membrane lipoprotein NlpD. Mutation of either *amiC* or *nlpD* in *N. meningitidis* completely abolished PG-derived peptide release (Fig. 2). AmiC and NlpD were also necessary for generating the free dipeptide. The enzyme responsible for dipeptide production has not been characterized in *Neisseria* spp., but three NlpC family endopeptidases are encoded in the meningococcal genome and these enzymes generally function to cleave the gamma-D-Glu-DAP bond [45]. The requirement for AmiC and NlpD for making the dipeptide suggests that the tetrapeptide must first be cleaved from the glycan strand by AmiC before the endopeptidase can act on it. In support of this idea, some NlpC family enzymes have been found to require a free N-terminal L-alanine for peptides to serve as substrates for these enzymes [33].

Mutation of *amiC* or *nlpD* led to a significant reduction in NOD1 activation. This result is consistent with the loss of free tripeptide release in these mutants. It is well established that human NOD1 responds to PG fragments containing gamma-D-Glu-DAP and that terminate with DAP, which would include the free tripeptide released by meningococci [18-20, 22]. In studies of *N. gonorrhoeae*, neither abolishing free peptide release nor abolishing disaccharide-tripeptide monomer release reduced NOD1 activation, indicating that production of either the tripeptides or the disaccharide-tripeptide monomers was sufficient for stimulating NOD1 [27]. Our results with *N. meningitidis* indicate that the free tripeptides are the major NOD1 agonists for this species and emphasize that meningococci rely on the AmiC-mediated pathway for most PG degradation (Fig. 1 and 2). Meningococci release lower levels of tripeptide PG monomer compared to gonococci, and meningococcal supernatant induces lower NOD1 activation in HEK293 cells and IL-8 production by Fallopian tube explants compared to gonococcal supernatant [8]. Taken together with these published observations, our results provide support to the hypothesis that *N. meningitidis* degrades PG fragments more extensively compared to *N. gonorrhoeae* and thereby reduces NOD1 activation, which may reduce immune clearance during asymptomatic colonization.

It is not currently clear how PG fragment release contributes to meningococcal lifestyle. In response to a NOD1 agonist or other inflammatory molecules, oral epithelial cells alter gene expression without inducing an inflammatory response [46]. Still, it is not particularly surprising that PG fragments released by meningococci do not induce inflammation in nasopharyngeal tissue, since asymptomatic colonization is by definition non-overtly damaging to the host on the organismal level. Released PG fragments may play a role in the establishment of colonization, in the invasion of the meninges or in exacerbating damage during invasive meningococcal disease, or in colonization of new niches such as the urethra (11, 55).

Consistent with the role of AmiC as a cell separation amidase, mutation of *amiC* or *nlpD* led to cell separation defects, which manifested as large cell clusters containing more than 10 cells (Fig. 4). STORM analyses revealed that in a small subset of cells, AmiC:FLAG3 formed a banding pattern at the septum, suggesting septal localization of this protein. In a larger subset of cells, AmiC:FLAG3 was found randomly distributed around the cell. NlpD:FLAG3 signal was found to localize to the septum. Such localization patterns suggest that NlpD is a septal protein and that AmiC is recruited to the septum during cell division, as observed in other species of bacteria [38, 44, 47].

The growth of the *amiC* mutant and the *nlpD* mutant as large aggregates of cells led us to test whether the cell separation deficiency might alter infection proficiency. In other systems, bacterial variants that grow as filaments have been found to have a survival advantage over normal bacilli and to avoid phagocytic killing [48]. However, gonococci defective for *amiC*, while not showing a growth defect, were sensitive to killing by deoxycholate, suggesting membrane instability [29]. Also, a meningococcal mutant lacking *gna33* (*ltgC*) showed a cell separation defect and was unable to infect infant rats in a model of septicemia [31]. We used a human whole blood infection model and infected the blood with the WT strain or an *nlpD* mutant carrying an inducible *nlpD* expression construct. While the WT strain was able to survive and grow in blood, the *nlpD* mutant showed more than a thousand fold defect in survival over 4h. The survival defect was still present, though substantially lessened, when blood depleted for complement was used for infections.

Overall, these studies demonstrated that *N. meningitidis* AmiC and NlpD are necessary for the producing the free peptides released by the bacteria and that meningococci favor this pathway for cell wall degradation. AmiC and NlpD are necessary for cell separation, and a cell separation mutant is defective in survival in an ex vivo model of human blood infection.

## MATERIALS AND METHODS

### Bacterial strains and growth conditions

All strains used in this study are listed in Table 1. Except where indicated, all meningococcal strains used in the experiments are unencapsulated derivatives of *N. meningitidis* (ATCC 13102 *cap*^-^), in which the capsule biosynthesis gene *siaD* has been interrupted with a chloramphenicol marker [8]. These strains also have a point mutation in *rpsL* conferring streptomycin resistance [8]. *N. meningitidis* strains were grown on gonococcal base medium (GCB) agar (Difco) plates at 37°C with 5% CO_2_ or in gonococcal base liquid medium (Difco) with 0.042% bicarbonate [49] and Kellogg’s supplements [50] (cGCBL) at 37°C with aeration. *Escherichia coli* strains were grown on LB agar (Difco) plates at 37°C or in LB liquid medium at 37°C. When needed, 10 μg/ml chloramphenicol, 10 μg/ml erythromycin, or 80 μg/ml kanamycin was added to *N. meningitidis* cultures. For *E. coli*, 100 μg/ml ampicillin, 25 μg/ml chloramphenicol, 500 μg/ml erythromycin, or 40 μg/ml kanamycin was added to the media when appropriate. When necessary, 0.1 mM or 1 mM IPTG, or 2 ng/ml anhydrotetracycline was added to the media to induce expression of a target gene.

**Table 1.**
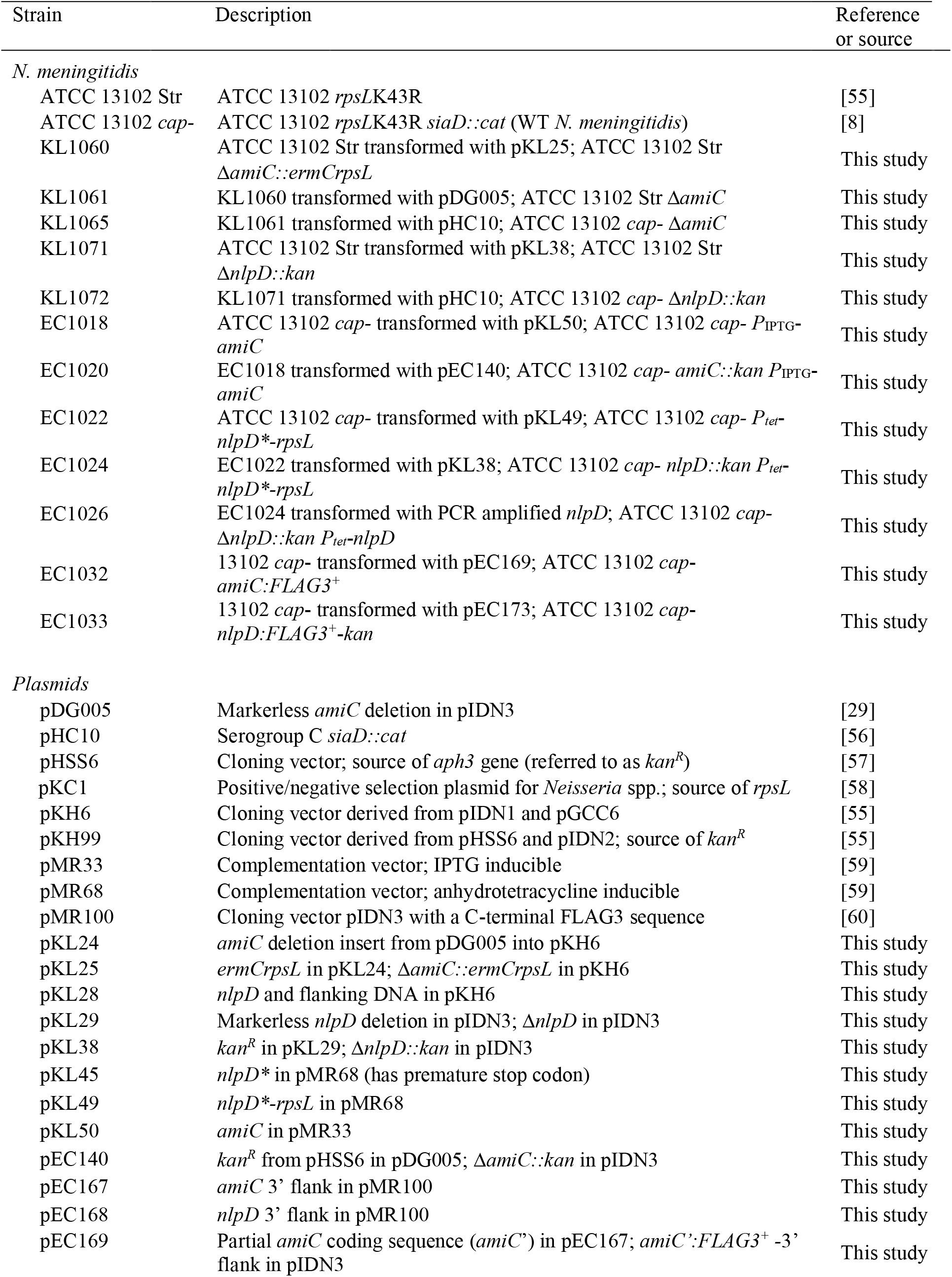
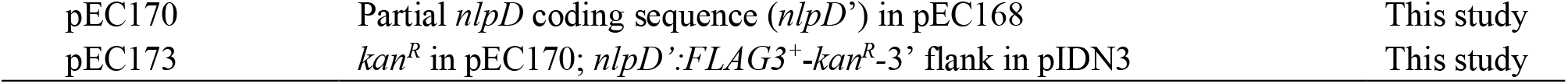
Strains used in this study.

### Strain construction

*N. meningitidis* mutants were generated by spot transformation [51]. Briefly, 20 μg linearized plasmid DNA or PCR product was spotted onto a GCB agar plate and allowed to dry. Subsequently, 5-10 piliated colonies were streaked over the DNA spots, and the plate was incubated overnight at 37°C with 5% CO_2_. Successful transformants were either selected using antibiotics, or identified by PCR screening. All transformants were confirmed by PCR and DNA sequencing. Due to the difficulty of transforming *amiC* and *nlpD* mutants, complementation strains were constructed by first transforming the WT with the complementation construct, followed by transformation of the resulting strains with the deletion construct. EC1026 is the only strain generated by transformation with a PCR product. This PCR product was generated by PCR amplification of *nlpD* from WT *N. meningitidis* chromosomal DNA using primer NlpD DUS F and NlpD DUS R (Table 2), which resulted in a DNA fragment containing the *nlpD* coding sequence flanked by DNA uptake sequences (DUS) to facilitate uptake and recombination [52]. Transformation with this PCR product corrected the *nlpD* sequence at the complementation locus to the WT sequence.

**Table 2.**
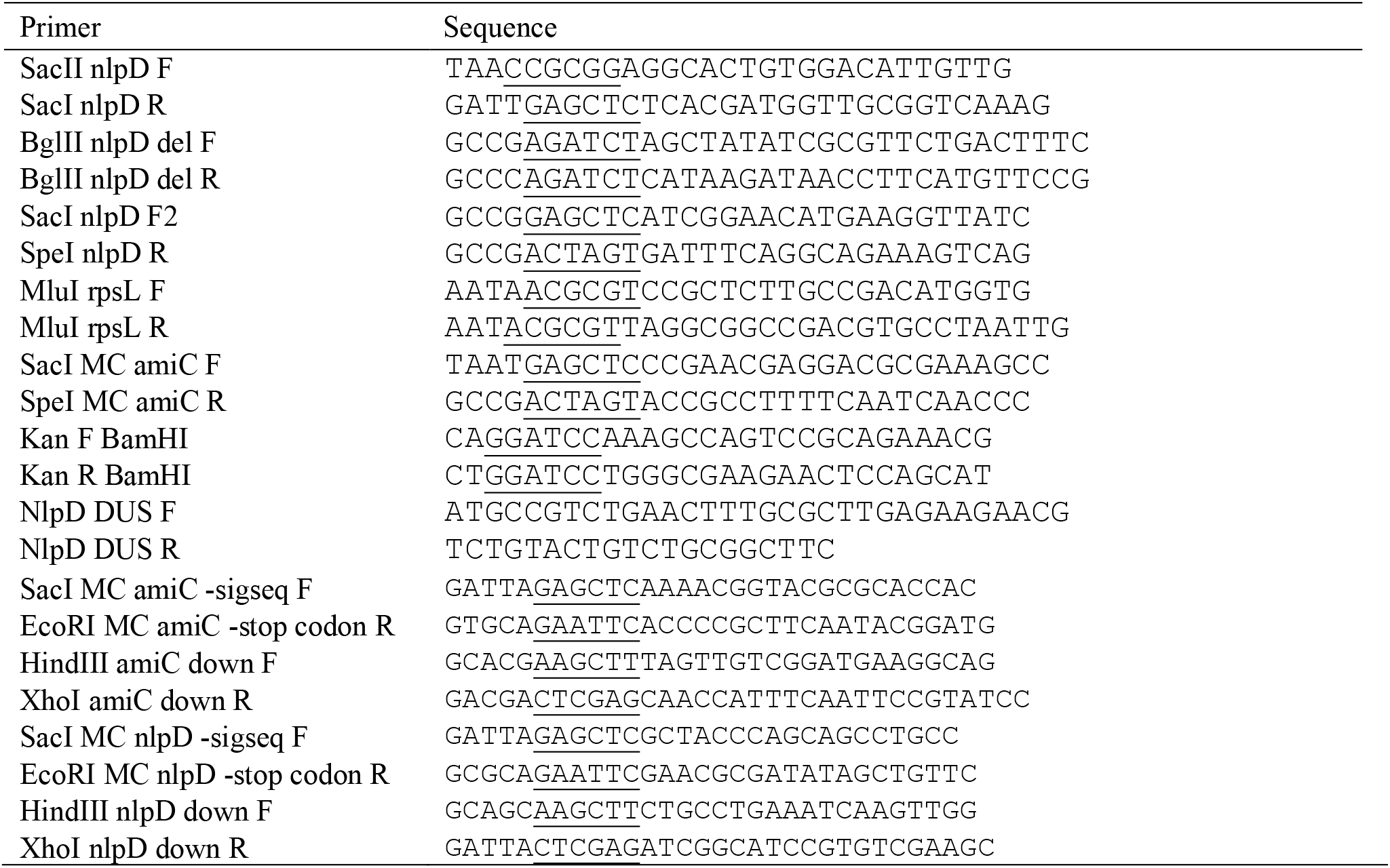
Primers used in this study. Restriction enzyme sites are underlined.

### Plasmid construction

All plasmids used in this study are listed in Table 1 and were verified by PCR and sequencing. All primers are listed in Table 2. Plasmids were generated by ligation of a vector backbone and an insert, and transformed into Rapid Trans™ chemically competent TAM1 cells (Active Motif).

Two plasmids, pKL24 and pKL25, were generated to create an in-frame deletion of *amiC*. To make pKL24 (*amiC* deletion in pKH6), pDG005 and pKH6 were digested with XmaI and SacI. The *amiC* deletion insert from pDG005 was ligated into pKH6 generating pKL24. To generate pKL25 (*ermC rpsL* in pKL24; final construct Δ*amiC::ermCrpsL*), the *ermC rpsL* positive/negative selection cassette was first excised from pKC1 by digestion with NheI and EcoRV. The insert from pKC1 was blunted with T4 polymerase and ligated into the blunted BspHI site from pKL24 to create pKL25.

Three plasmids were generated to create a *nlpD* insertional inactivation mutant. To create pKL28 (*nlpD* and flanking DNA in pKH6), PCR was used to amplify *nlpD* and flanking DNA with primers SacII nlpD F and SacI nlpD R. The PCR product and pKH6 were digested with SacI and SacII, then ligated. To generate pKL29 (*nlpD* deletion in pKH6), primers BglII nlpD del F and BglII nlpD del R were used in a PCR with pKL28 as a template. The product was digested with BglII and self-ligated to make pKL29, which contains a markerless *nlpD* deletion with flanking DNA. To build pKL38 (Δ*nlpD::kan* allelic replacement in pKH6), the kanamycin resistance marker from pKH99 was excised by digestion with EcoRV and Ecl136II and inserted into the blunted BglII site of pKL29.

Two plasmids, pKL45 and pKL49, were constructed to complement an *nlpD* mutation at an ectopic site. Plasmid pKL45 (*nlpD\** in pMR68) was made by first amplifying *nlpD* from WT *N. meningitidis* chromosomal DNA with primers SacI nlpD F2 and SpeI nlpD R. The PCR product and pMR68 were subsequently digested with SacI and SpeI and ligated to each other. Despite repeated attempts, the resulting plasmid contained one or more mutations in the *nlpD* coding sequence. We deduce that the presence of a second copy of *nlpD* in *E. coli* is toxic to the bacterium, and saved one of the plasmids as pKL45. pKL45 has a nonsense mutation leading to a premature stop codon at the third codon. Plasmid pKL49 (*nlpD*-rpsL* in pMR68) was generated by insertion of *rpsL* into pKL48. *rpsL* was amplified from pKC1 using primers MluI rspL F and MluI rpsL R; the PCR product and pKL45 were digested with MluI and subsequently ligated to each other to generate pKL49.

Plasmid pKL50 was generated to complement an *amiC* mutation at an ectopic site. To make pKL50 (*amiC* in pMR33), *amiC* was amplified from WT *N. meningitidis* chromosomal DNA with primers SacI MC amiC F and SpeI MC amiC R and digested with SacI and SpeI. pMR33 was digested similarly, and ligated with the digested *amiC* PCR product.

Plasmid pEC140 was built to create a deletion/insertion mutant where *kan* replaced *amiC*. To generate pEC140 (*amiC::kan* in pIDN3), pDG005 was first digested with BspHI and subsequently blunted with T4 DNA polymerase. The *aph3* gene conferring resistance against kanamycin (referred to as *kan*^*R*^) was amplified from pHSS6 using primers kan F BamHI and kan R BamHI and blunt-ligated with the blunted, BspHI digested pDG005 to form pEC140.

To epitope tag *amiC* at the native locus, pEC169 (*amiC’:FLAG3*^*+*^ *-*3’ flank in pIDN3) was constructed in multiple steps. First, *amiC* 3’ flank region was amplified using primers HindIII MC amiC down F and XhoI MC amiC down R and WT *N. meningitidis* chromosomal DNA as a template. pMR100 and the PCR product were digested with HindIII and XhoI, and ligated to make pEC167 (*amiC* 3’ flank in pMR100). The partial coding sequence of *amiC* lacking the start codon, signal sequence and stop codon (*amiC’*) was amplified from WT *N. meningitidis* DNA using primers SacI MC amiC –sigseq F and EcoRI MC amiC –stop codon R, digested with SacI and EcoRI and ligated into similarly digested pEC167 to form pEC169.

To epitope tag *nlpD* at the native locus, pEC173 (*nlpD’:FLAG3*^*+*^*-kan*^*R*^*-*3’ flank in pIDN3) was generated with a similar strategy as pEC169. *nlpD* 3’ flank region was amplified using primers HindIII MC nlpD down F and XhoI MC nlpD down R instead, digested with HindIII and XhoI and ligated with similarly digested pMR100 to form pEC168. The partial coding sequence of *nlpD* lacking the start codon, signal sequence and stop codon (*nlpD’*) was amplified with primers SacI MC nlpD –sigseq F and EcoRI MC nlpD –stop codon R, digested with SacI and EcoRI and ligated into similarly digested pEC168 to form pEC170. pEC170 was digested with HindIII, blunted with T4 polymerase and ligated with *kan*^*R*^ to form pEC173.

### Metabolic labeling of peptidoglycan and quantitative fragment release

Metabolic labeling of PG and quantitative fragment release was performed as described previously [53]. Strains were pulse-chased labeled with 25 μg/ml [2,6-^3^H] DAP in DMEM lacking cysteine supplemented with 100 µg/ml methionine and 100 µg/ml threonine for 45 minutes. A sample of the culture was removed to determine the number of radioactive counts per minute (CPM) by liquid scintillation counting, and the bacteria were diluted to obtain equal CPM in each culture. After a 2 hour chase period, supernatant was harvested by centrifuging the culture at 3,000 x *g* for 10 mins, and filter-sterilizing the supernatant with a 0.22 μm filter. [2,6-^3^H] DAP labeled molecules in the filtered supernatant were then separated using tandem size-exclusion chromatography with 0.1 M LiCl as the mobile phase, and detected by liquid scintillation counting. The growth medium for the *amiC* and *nlpD* complementation strains was supplemented with 0.1 mM IPTG and 2 ng/ml anhydrotetracyline, respectively.

### Thin section transmission electron microscopy

*N. meningitidis* strains were grown in cGCBL until mid-log phase, when cells were harvested by centrifugation at 17, 000 x *g* for 1 minute. Growth medium for the *amiC* and *nlpD* complementation strains contained 0.1 mM IPTG or 2 ng/ml anhydrotetracycline, respectively, for gene induction. Cells were washed once with PBS and resuspended in fixative solution (2% paraformaldehye, 2.5% glutaraldehye in 0.1M phosphate buffer). Thin section electron microscopy of fixed samples was done at the University of Wisconsin-Madison Medical School Electron Microscope Facility. Enumeration of meningococci in cell clusters were done by counting the number of complete, in plane cells presenting as monococci, diplococci, tetrads or clusters of five or more cells in 16-24 fields/strain containing an average of 30 cells/field.

### Treatment of HEK293 reporter cell line with filtered supernatant from meningococcal cultures

HEK293-Blue cells that encode secreted alkaline phosphatase (SEAP) under the control of an NF-κB promoter (parental cell lines NULL1 and NULL2) and hNOD1 or hNOD2 receptors (NOD and NOD2 cell lines) were grown, maintained and treated according to the manufacturer’s instructions (InvivoGen). Supernatant samples were harvested from meningococcal cultures started at OD_540_ of 0.2 and grown for 2 hours by centrifugation at 17,000 x *g* for 1 minute and filter sterilized using a 0.22 µm pore filter. Cultures were supplemented with 1 mM IPTG or 2 ng/ml anhydrotetracycline as needed. Supernatant samples were normalized to total cellular protein. HEK293 cells that were seeded at a density of 2.8 × 10^5^ cells/ml (NOD1/NULL1) or 1.4 × 10^5^ cells/ml (NOD2/NULL2) in 180 µl DMEM with 4.5g/L glucose, 2 mM L-glutamine and 10% heat inactivated fetal bovine serum (FBS) were treated with 20 µl supernatant for 18 hours at 37°C with 5% CO_2_. A total volume of 20 µl each of cGCBL (media only control), 10 µg/ml TriDAP (positive control for NOD1/NULL1) (InvivoGen) or 10 µg/ml muramyl dipeptide (MDP; positive control for NOD2/NULL2) (InvivoGen), 10 µg/ml Tri-Lys (negative control for NOD1/NULL1) or 10 µg/ml MDP-control (MDP-c; negative control for NOD2/NULL2) (InvivoGen) were used as treatment controls. After 18 hours of treatment, 20 µl HEK293 culture supernatant was added to 180 µl QUANTI-Blue reagent (InvivoGen) and incubated at 37°C for 4 hours. SEAP activity was determined by measuring absorbance at 650 nm. To calculate NF-κB dependent hNOD activation (graphed SEAP levels (A_650_)), A_650_ values from NULL1 or NULL2 controls were subtracted from the corresponding A_650_ values from NOD1 or NOD2 treatments. Results are from four independent experiments, and statistical tests were performed using Student’s *t-*test.

### Stochastic optical reconstruction microscopy (STORM)

Sample preparation and imaging was done as described previously [23]. Meningococcal cultures were grown to mid-log phase and harvested by centrifugation at 17,000 x *g* for 2 minutes. Cells were washed once with PBS, and suspended in 400 µl fixative solution (4% methanol-free formaldehyde in PBS). The cell suspensions were first incubated at RT for 15 minutes, and then on ice for 15 minutes. The cell suspensions were then centrifuged at 6,000 x *g* and the fixative solution discarded. Cells were quickly washed thrice with cold PBS to ensure thorough removal of fixative solution, and resuspended in cold GTE buffer (50 mM glucose, 1mM EDTA, 20 mM Tris-HCl pH 7.5) and incubated on ice for 10 minutes. Cells were quickly washed with PBS-T (0.3% Triton-X100 in PBS), suspended in cold methanol and incubated at -20°C for a minimum of 10 minutes. Methanol was removed, after which cells were washed with PBS-T for 5 minutes, blocked with 400 µl 5% goat serum in PBS-T for 15 minutes, and incubated with 1:150 diluted M2 primary antibody (Sigma) in PBS for 1 hour. Cells were subsequently washed with PBS thrice, and incubated with 200 µl 1:100 diluted AlexaFluor-647 (Thermo Fisher) secondary antibody, 100 ng/ml DAPI and 5% goat serum in PBS for 1 hour at RT or O/N at 4°C. Cells were washed with PBS five times and suspended in 200 µl PBS to remove unbound antibody and dye. Around 2 µl volumes of the samples were spotted onto poly-L-lysine coated coverslips, mounted with VECTASHIELD™ H-1000 mounting media (Vector laboratories) and visualized using a Nikon N-STORM microscope. Images were analyzed using the software NIS-Elements AR with N-STORM analysis. Linear brightness/contrast adjustments and false-coloring was performed using FIJI.

### Human blood infection assay

Blood was collected from healthy donors using hirudin-containing vacutainer tubes [54]. All human subjects gave written informed consent in accordance with a protocol approved by the University of Wisconsin-Madison Health Sciences Institutional Review Board (Protocol #2017-1179). *N. meningitidis* was grown overnight on a GCB agar plate and harvested into phosphate-buffered saline. The blood was inoculated with *N. meningitidis* strains at 1 × 10^8^ CFU/ml bacteria, and the cultures were grown with rotation at 37°C. To enumerate *N. meningitidis* used as the inoculum at the start of the experiment, the bacteria were inoculated into GCBL instead of blood and dilution plated. By contrast, t=0 was the time point taken immediately after inoculation of the blood. Where indicated, blood was pretreated with 1µg/ml cobra venom factor for 1 hour prior to inoculation, with the blood culture rotating at 37°C. At each time point 375µl of each blood culture was removed and diluted into 1.125ml of cGCBL medium. Anhydrotetracycline (final concentration 2ng/ml) was added to each culture. The diluted cultures were grown at 37°C for 4 hours to allow for expression from the complementation construct. After 4 hours, the bacteria were diluted and plated for CFU determinations.

## ACKNOWLEDGEMENTS

This work was supported by the National Institute of Allergy and Infectious Disease grant R01AI097157 to J.P.D. We would like to acknowledge Ben August from the UW-Madison Medical School EM Facility for assistance with TEM, and Elle Kielar-Grevstad from the UW-Madison Biochemistry Optical Core for advice and technical assistance on STORM.

